# Differential expression of proteins in human prostate cancer tissues probed by MALDI imaging mass spectrometry

**DOI:** 10.1101/2020.10.05.326686

**Authors:** Amrita Mitra, Surya Kant Choubey, Rajdeep Das, Pritilata Rout, T. S Sridhar, Deepak Mishra, Amit Kumar Mandal

**Author notes:** Corresponding Author, Dr. Amit Kumar Mandal, Clinical Proteomics Unit, Division of Molecular Medicine, St. John’s Research Institute, St. John’s National Academy of Health Sciences, 100ft Road, Koramangala, Bangalore – 560034, India. Dr. Amit Kumar Mandal, Department of Biological Sciences, Indian Institute of Science Education and Research Kolkata, Mohanpur, Nadia, West Bengal, India, Pin-741246.

## Abstract

Histopathology, the gold-standard method for diagnosis of human prostate cancer, is based on the analysis of changes in cellular morphology and tissue architecture in chemically stained tissue sections. Even at the very early stages of cancer with minimal phenotypic changes in cellular morphologies, there might be detectable changes in the expression profiles of proteins. Over the last decade, imaging mass spectrometry has been used to explore the spatial distribution and expression profiles of several molecules with their twodimensional heterogeneity retained across the tissue section. In the present study, using MALDI mass spectrometry based tissue imaging, we report the differential expression of three proteins, vinculin, ribonuclease T2 and 60 kDa heat shock protein across human prostate tissue sections in regions pathologically demarcated as frankly malignant. In an independent analysis, quantitative proteomics revealed that in cancerous prostate tissues, ribonuclease T2 and 60 kDa heat shock protein were significantly overexpressed by 22.26- and 6 folds respectively compared to benign condition. Our results show the utility of this approach to probe differential protein expression in architecturally intact cancer tissues. In addition, we propose that ribonuclease T2 and 60 kDa heat shock protein might be developed as diagnostic biomarkers for prostate cancer in future.

## Introduction

Prostate cancer is one of the most common type of cancer in men over the age of 60 years [1,2]. Although a significant proportion of prostate cancer initially tends to grow slowly, they ultimately develop into lesions with clinical manifestation and metastasize [3]. In the routine diagnosis of prostate cancer, prostate specific antigen (PSA), a peripheral blood biomarker, is quantitatively assayed to detect prostate cancer with a cut-off limit of <4 ng/mL of PSA in blood under normal conditions [4]. It has been reported that in several cases, prevalence of prostate cancer was observed in patients even when the PSA levels were below 4 ng/mL [5]. Thus, PSA evaluation is not a reliable method for the diagnosis of prostate cancer. In this scenario, the gold standard approach for the definitive diagnosis of prostate cancer is based on the histopathological examination of tissue biopsies. However, it has been observed that histopathological findings might be subject to the inter-pathologist discordance over the Gleason scoring of tissues, where 12.7% needle core biopsies are reportedly overgraded and 50.1% are undergraded [6,7]. In addition, histopathological examination of prostate needle core biopsies has been reported with a false negative rate in the detection of cancer by approximately 2.4% [8]. Thus, for the unambiguous diagnosis of the cancer with high precision and also to gain molecular insights of the disease, a molecular profile based diagnostic approach might be crucial.

Over the last decade, Matrix-Assisted Laser Desorption/Ionization (MALDI) mass spectrometry based tissue imaging and Desorption Electrospray Ionization (DESI) mass spectrometry based tissue imaging have been used to visualize the spatial distribution of several proteins and metabolites in a single scan without the perturbation of the two-dimensional heterogeneity of their distribution across a tissue section [9–11]. In imaging mass spectrometry (IMS), for a given mass-to-charge (m/z) ratio the molecular images are constructed by using a cartesian coordinate system, where the data are represented by the x and y coordinates and signal intensity (i) is proportional to the abundance of the corresponding molecular ion. The variation in the intensity of each molecular ion across the tissue section is visualized is visualized as a heat map of ion intensity relative to a color scale. A full imaging MS data consists of the information in a 2+n dimensional space (x/y/i_n_), where n is the number of m/z signals detected during data acquisition. An intensity plot or image is constructed for each m/z value [12].

In the present study, MALDI mass spectrometry based tissue imaging was used to visualize the distribution of various proteins across human prostate tissues which were assigned as cancerous upon histopathological examination. Using nLC/ESI-MS based tissue proteomics approach, we also identified, characterised and quantified the differential expression profiles of these proteins in the cancerous prostate tissues and compared the same with that in the benign prostate tissue.

## Materials and methods

### Materials

Superfrost™ microscopic slides were purchased from Thermo Fisher (USA). Analytical grade ethanol, glacial acetic acid, chloroform and red phosphorous were obtained from Merck (Germany). Sequencing grade bovine trypsin, β-octyl glucopyranoside and α-Cyano-4-hydroxycinnamic acid matrix were purchased from Sigma Aldrich (St. Louis, MO). Water and acetonitrile were of LC–MS grade and obtained from Honeywell (California, USA). All other chemicals used were of analytical grade.

### Ethics statement

The study was approved by the Institutional Ethical Committee (IEC approval no. 27/2014), St. John’s Medical College, Bangalore, Karnataka, India. All the procedures were carried out in accordance with the available guidelines and regulations for human participants in research.

### Selection of subjects

Patients presenting with prostate cancer and benign prostatic hyperplasia (BPH) to the Department of Urology at St. John’s Medical College Hospital, Bangalore, were invited to participate in the study. Written consents were obtained from patients prior to collection of samples.

### Collection and sectioning of prostate tissue samples

Twenty needle core biopsies were collected in normal saline from ten patients suspected with prostate cancer, in addition to one prostate tissue core that was obtained from a patient undergoing the procedure of transurethral resection of the prostate (TURP). Out of the collected needle core biopsies, seven were diagnosed with prostate cancer using routine histopathological examination. Additionally, twenty tissue cores from ten patients with BPH undergoing the procedure of TURP were also collected. The tissue samples were mounted on microtome chucks and fixed in ice for sectioning in a cryomicrotome. Sections of 8 μm thickness were taken and thaw-mounted on microscopic glass slides. Two serial sections for each tissue were collected, one each for histopathological examination and MALDI imaging experiment. The remaining cores of tissues were desiccated and stored in −80 °C until used for tissue proteomics experiments.

### Processing of tissue section for MALDI imaging experiment

The slides containing the tissue sections were desiccated for 45 mins. For fixing the tissue sections and removal of lipids and salts, the tissue sections were treated stepwise with different solutions as follows: 70% ethanol (30 seconds), 100% ethanol (30 seconds), 6:3:1 of ethanol: chloroform: glacial acetic acid – Carnoy’s solution (2 minutes), 100% ethanol (30 seconds), 0.2% trifluoroacetic acid in water (30 seconds) and 100% ethanol (30 seconds). Subsequently, the slides with tissue sections were desiccated for 30 minutes to remove all the solvents, followed by *in situ* trypsin digestion and spraying of matrix.

### *In situ* trypsin digestion and matrix application

Trypsin solution (0.1 mg/ml) in 50 mM NH_4_HCO_3_, pH 7.4 containing 1% β-octylglucopyranoside was sprayed over a tissue section at the flow rate of 6 μl/min for 10 layers (Suncollect sprayer/spotter, Germany). The slides were desiccated for 30 minutes and incubated at 37 °C overnight. α-Cyano-4-hydroxycinnamic acid (HCCA) (5 mg/ml in 1:1 ACN/0.5% aqueous TFA) matrix was sprayed over the tissue sections at the flow rate of 10 μl/min for the first two layers, followed by 15 μl/min for the next 6 cycles and 20 μl/min for the last two cycles.

### MALDI MS imaging of tissue sections

The samples were analyzed using a MALDI Q-TOF mass spectrometer calibrated with red phosphorous (Synapt G2S*i*, WATERS, UK). Mass spectra were acquired in positive polarity in the m/z range 1000 to 4500 with a scan time of 1 sec using the analyzer in sensitivity mode. In general, the quadrupole mass analyzer that was used here in the MS scan, does not transmit all ions with the equal efficiency. Rather, it acts as a broad-band filter that allows passage of ions with m/z values from 0.8 × mass 1 (lower mass end). On the high mass end, the transmission of ions occurs for m/z up to 5 × mass 3. To allow equal transmission of ions in the m/z range 1000 to 4500, the following quadrupole ion transmission profile used: mass 1 (m/z 100), dwell time (4%), ramp time from mass 1 to mass 2 (1%), mass 2 (m/z 900), dwell time (90%), ramp time from mass 2 to mass 3 (5%), and mass 3 (m/z 900). The spatial resolution, measured in terms of pixel size, was optimized at 90 μm in both X and Y directions. The laser was operated at a repetition rate of 1000 Hz and with laser energy of 370 kV. The instrument was operated using a trap collision energy of 10 kV and trap cooling nitrogen gas flow rate of 2.5 mL/min. Image acquisition was carried out using High Definition Imaging (HDI) software, v 1.4 coupled to MassLynx v 4.1 (WATERS, UK).

### Analysis of images using HDI v 1.4 software

The acquired data were analyzed using HDI software (WATERS, UK). One thousand most intense peaks inclusive of their isotopic distributions were selected from the acquired data for processing in HDI software and to match their specific localization on the tissue sections in terms of X and Y coordinates. The parameters used for processing the data were as follows: m/z window = 0.1 Da; MS resolution = 12000; internal lock mass peak = 2272.1; lock mass tolerance = 0.3 amu; lock mass minimum signal intensity = 500 counts. The intensities of all the peaks were normalized by the total ion count (TIC), where the absolute intensities of each peak was divided by the sum of the absolute intensities of all peaks. The normalized values were plotted to obtain the images corresponding to the distribution of multiple peptides on the tissue sections.

### Tissue proteomics of human prostate samples

#### Processing of the collected tissue samples

One core of prostate tissue samples was lyophilized for 4 hours. The lyophilized tissue was crushed to powder. 0.2 mg of the crushed sample was suspended in tissue lysis buffer containing 10 mM tris HCl, 0.15 M NaCl, 1 mM EDTA in phosphate buffered saline, pH 7.4 with 1% β-octyl glucopyranoside. 10 μL of protease inhibitor cocktail was added to the suspension and subsequently sonicated for 30 min with intermittent cooling. After sonication, the solution was incubated for 1 hour at 4 °C with mild agitation. The suspension was centrifuged at 14,700 x g for 45 min and the supernatant was dialyzed overnight with frequent changes against 50 mM NH4HCO3 buffer, pH 7.4. After dialysis, the protein concentration was measured by Bradford assay method according to the manufacturer’s protocol and followed by in-solution digestion.

### In-solution digestion of the proteins from tissue samples

Samples were denatured by using 0.2% Rapigest at 80 °C for 15 min. Subsequently, the disulphide linkages of proteins were reduced by 5 mM dithiothreitol for 30 min at 60 °C, followed by alkylation of the reduced –SH groups with 10 mM iodoacetamide for 30 min in dark at room temperature. Modified sequencing grade trypsin, reconstituted in 50 mM NH_4_HCO_3_, pH 7.4, was added to the protein extracts at an enzyme:substrate ratio of 1:10 and incubated overnight at 37 °C. Following this, the samples were acidified in 0.5% (v/v) formic acid (FA) and incubated at 37 °C for 90 mins to facilitate hydrolysis of Rapigest detergent. Samples were then be centrifuged for 45 min at 4000 x g and the supernatant was collected and spiked with 50 fmoles/μl of internal standard yeast enolase (WATERS, UK). Subsequently, the samples were carried forward for nanoLC-ESI-MS analysis.

### nanoLC fractionation

For tissue proteomics, the samples were analyzed at Proteomics Facility, Centre for Cellular and Molecular Platforms (CCAMP), National Centre for Biological Sciences (NCBS), Bangalore. The proteolytic peptides were fractionated in EASY-nanoLC 1200 UHPLC system (Thermo Scientific, Germany), calibrated using Pierce™ LTQ ESI Positive Ion Calibration Solution (Thermo Scientific, Germany). Briefly, 600 ng of total tryptic digest of the experimental sample was loaded directly into a C18 Acclaim™ PepMap™ Rapid Separation Liquid Chromatography column (3 μm, 100 Å x 75 μm x 50 cm). The aqueous phase consisted of water/0.1% FA (solvent A) and the organic phase comprised of acetonitrile/0.1% FA (solvent B). The flow rate of solvents through the column was 300 nL/min. A 90 min run program was designed, starting with a linear gradient of solvent B from 5-40% for 70 min, followed by 40-80% of solvent B for 8 min, and finally 80-95% of solvent B for 12 min. The column temperature was maintained at 40 °C during the run in all experiments. The eluted peptides were analyzed in a mass spectrometer with an orbitrap mass analyser (Orbitrap Fusion™ mass spectrometer, Thermo Scientific, San Jose, California, USA).

### Data acquisition

The data were acquired in the mass range of 375-1700 m/z at 120000 FWHM resolution (at 200 m/z). The instrument was operated in positive ion mode with a source temperature of 350 °C, capillary voltage of 2.0 kV. Poly-siloxane (445.12003 m/z), used as lock mass, was continuously infused throughout the acquisition. The instrument was set to run in top speed mode with 3 s cycles for the survey and the MS/MS scans. After a survey scan, tandem MS was performed on the most abundant precursors exhibiting charge states from 2 to 7 with the intensity greater than 1.0e^4^, by isolating them in the quadrupole. Higher-energy collisional dissociation (HCD) was used for fragmentation of the ions where the normalized collision energy was maintained at 35% and resulting fragments were detected using the rapid scan rate in the ion trap. The automatic gain control (AGC) target for MS/MS was set to 4.0e^5^ and the maximum injection time limited to 50 msec. The experimental samples were run in duplicate.

### Proteomics Data Analysis

The acquired data was processed using Thermo Scientific™ Proteome Discoverer™ software version 2.1. MS/MS spectra were searched with the SEQUEST^®^ HT engine against human proteome database incorporated with yeast enolase sequence details. During analysis, the parameters used were as follows: proteolytic enzyme, trypsin; missed cleavages allowed, two; minimum peptide length, 6 amino acids. Carbamidomethylation (+57.021 Da) at cysteine residues was kept as a fixed modification. Methionine oxidation (+15.9949 Da) and protein N-terminal acetylation (+42.0106 Da) were set as variable modifications. The mass tolerances for precursor and product ions were kept at 10 ppm and 0.6 Da respectively. Peptides were confidently matched to protein sequences with a maximum false discovery rate of 1% as determined by the Percolator^®^ algorithm. Protein groups were further identified with a criteria of having the presence of at least two unique peptides.

## Results and discussion

The gold standard method for the definitive tissue based diagnosis of cancer is microscopy of stained tissue sections evaluated by a trained pathologist. In this procedure, tissue diagnosis and subsequent gradation of cancers are based on the characterization of cellular morphology and alteration of tissue architecture in hematoxylin and eosin (H & E) stained tissue sections [13]. Using MALDI mass spectrometry based tissue imaging platform, we imaged various proteins across seven prostate tissue sections which were diagnosed as cancerous based on histopathological examination. Three proteins, vinculin, ribonuclease T2 and 60 kDa heat shock protein were observed to be differentially distributed across the tissue sections. The differential distributions of these three proteins were consistently observed in the needle core biopsies of the prostate gland isolated from patients with prostate cancer. In addition, we performed an independent quantitative tissue proteomics study to investigate the expression of these proteins in tissue proteomes in prostate cancer in comparison to that in BPH. Proteomics data showed that ribonuclease T2 and 60 kDa heat shock protein were significantly overexpressed by 22.26 and 6 folds respectively.

Figure 1A represents the digital image of an H & E stained prostate cancer tissue section. Figures 1B, 1C, 1D and 1E represent the 20 X views of different regions of the tissue section showing distribution of tumor cells observed in variable amounts of stroma. Figures 1B and 1C show the lobular, nests and acinar patterns of the cancer cells, separated by fibrous stroma infiltrated with scattered lymphocytes. Figures 1D and 1E show the arrangement of cancerous cells as acini and glands.

**Figure 1:**
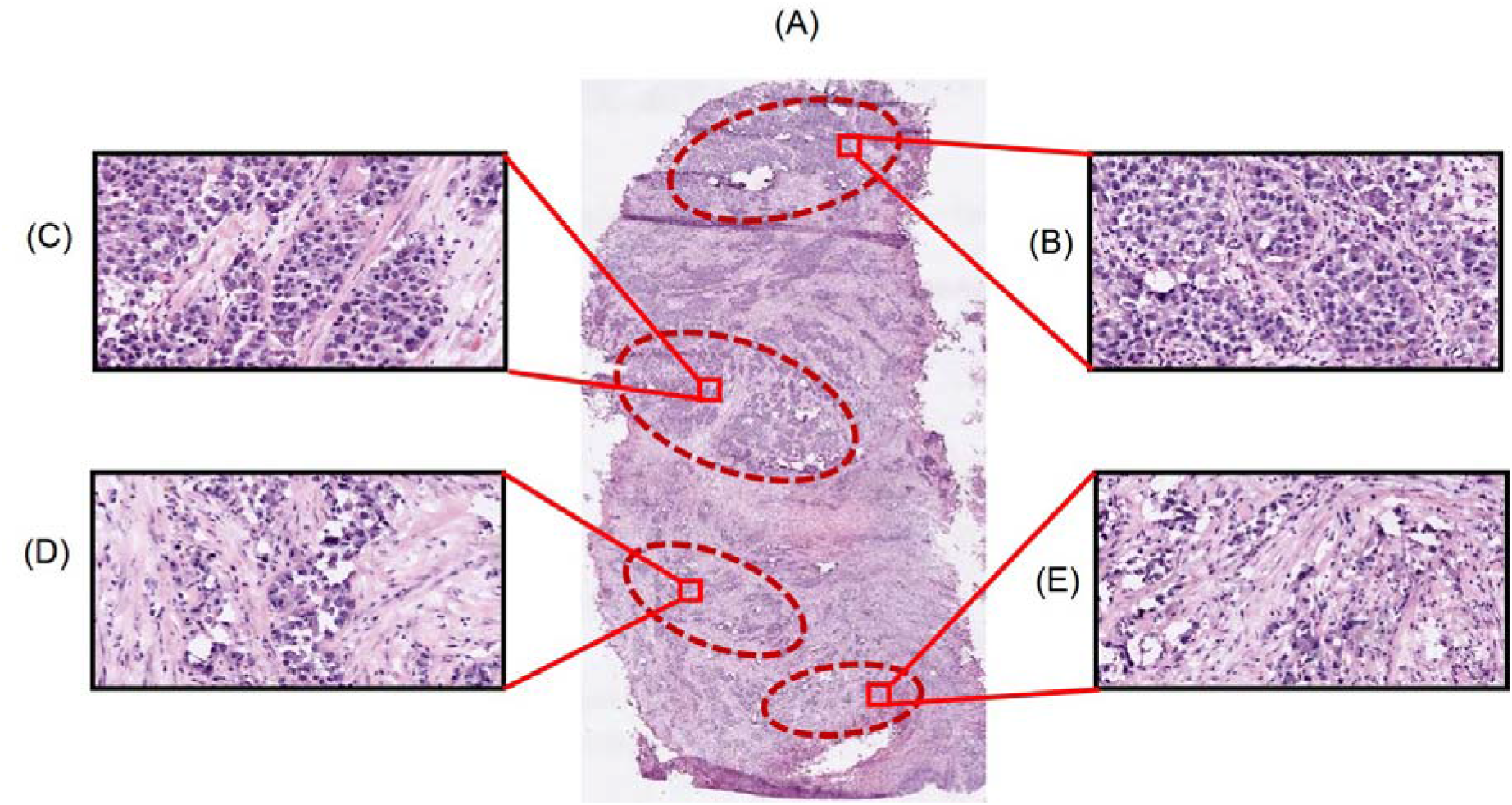
Representation of the cellular population in different regions of the human prostate cancer tissue section. Panel A shows the tissue section (11 mm × 6 mm) obtained from a patient with confirmed prostate cancer (Age: 68 yrs; Gleason’s grade 7 (3+4) poorly differentiated carcinoma of urothelial origin), stained with H & E and scanned with a magnification of 20X. Panels B, C, D and E represent the overall cellular population in different pathologically significant regions of the prostate tissue section

Figure 2 represents the combined mass spectra of various molecular ions distributed across the entire region of the tissue section. Ninety proteolytic peptides were observed in the mass spectra. Upon analysis of the nLC/ESI-MS based tissue proteomics data, fifty two proteins could be identified and characterized from these peptides. The intensities of these peptides as observed in the MALDI spectra were plotted along the X and Y coordinates and subsequently the images of the distribution of these peptides were obtained in the experimental tissue sections. In the MALDI images, forty-one proteins were observed with low sensitivity such that their differential spatial distribution was unclear. These proteins were discarded from the analysis. In addition, eight proteins were observed with a uniform pattern of distribution across the tissue section. These were characterized as fragments of trypsin, which was used for proteolytic digestion *in situ*, hemoglobin and of the different isoforms of cytoskeletal proteins, actin and myosin. The remaining three proteins, vinculin, ribonuclease T2 and 60 kDa heat shock protein were the proteins of our interest as their differential distribution across the tissue sections could be attributed to prostate cancer.

**Figure 2:**
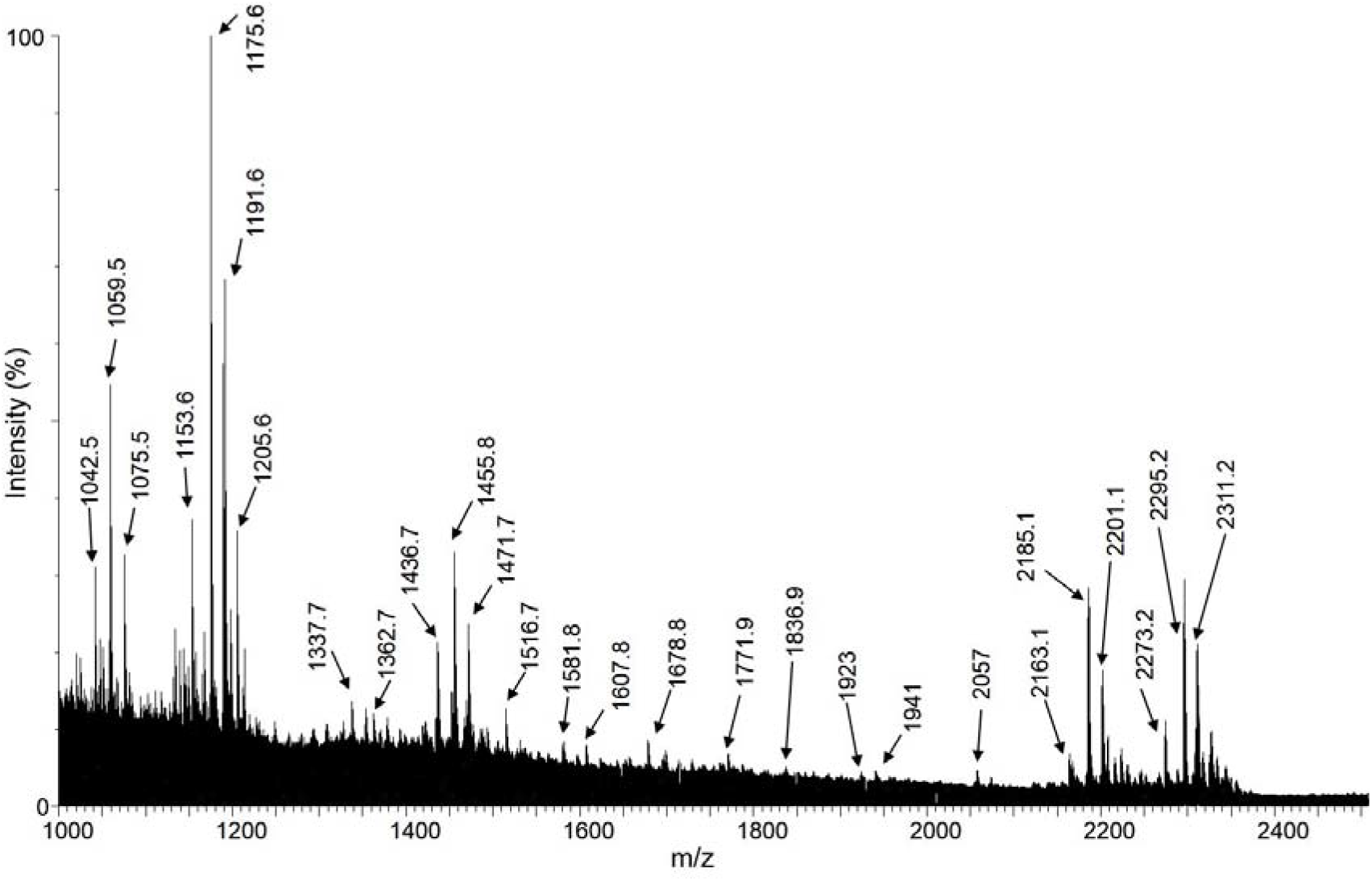
The combined MALDI mass spectra obtained from all pixel points within a prostate cancer tissue section

Figure 3 shows the homogeneous distribution of a peptide with m/z 1455.8 across the experimental prostate cancer tissue section in the mass spectrometry based MALDI image. nLC/ESI-MS based tissue proteomics identified this peptide as a tryptic fragment of the cytoskeletal protein myosin-11, spanning residues ^1365^HISTLNIQLSDSK^1377^. Figure 3A shows the corresponding representative regions in the H & E stained tissue section (encircled) where myosin-11 was observed as homogeneously expressed in the MALDI image of a serial section of the same tissue, depicted in Figure 3B. The isotopic distribution of the peptide with m/z 1455.8 in the mass spectra is shown in Figure 3C. Figures 3D and 3E represent the 20X expanded views in a small representative part of the encircled areas showing cellular populations in these regions. Upon analyzing the cellular population in the H & E stained tissue section, we observed that myosin-11 was homogeneously expressed in the regions having high density of cells as well as other regions having greater amounts of stroma with immune cells. It has been reported that myosin-11 is expressed in the smooth muscle cells of the prostate gland [14]. Its major function is in smooth muscle contraction upon association with actin microfilament proteins [15,16]. Additionally, in a prostate cancer tissue, the smooth muscle cells are also contributed by the new blood vessels formed during angiogenesis (the process of formation of new blood vessels through tumors), whose fine branches can spread across the entire tissue section [17]. In the present study, the uniform distribution of myosin-11 across the tissue section might owe to the abovementioned factors. Therefore, our observation correlates well with reported cytological features of human prostate cancer tissues.

**Figure 3:**
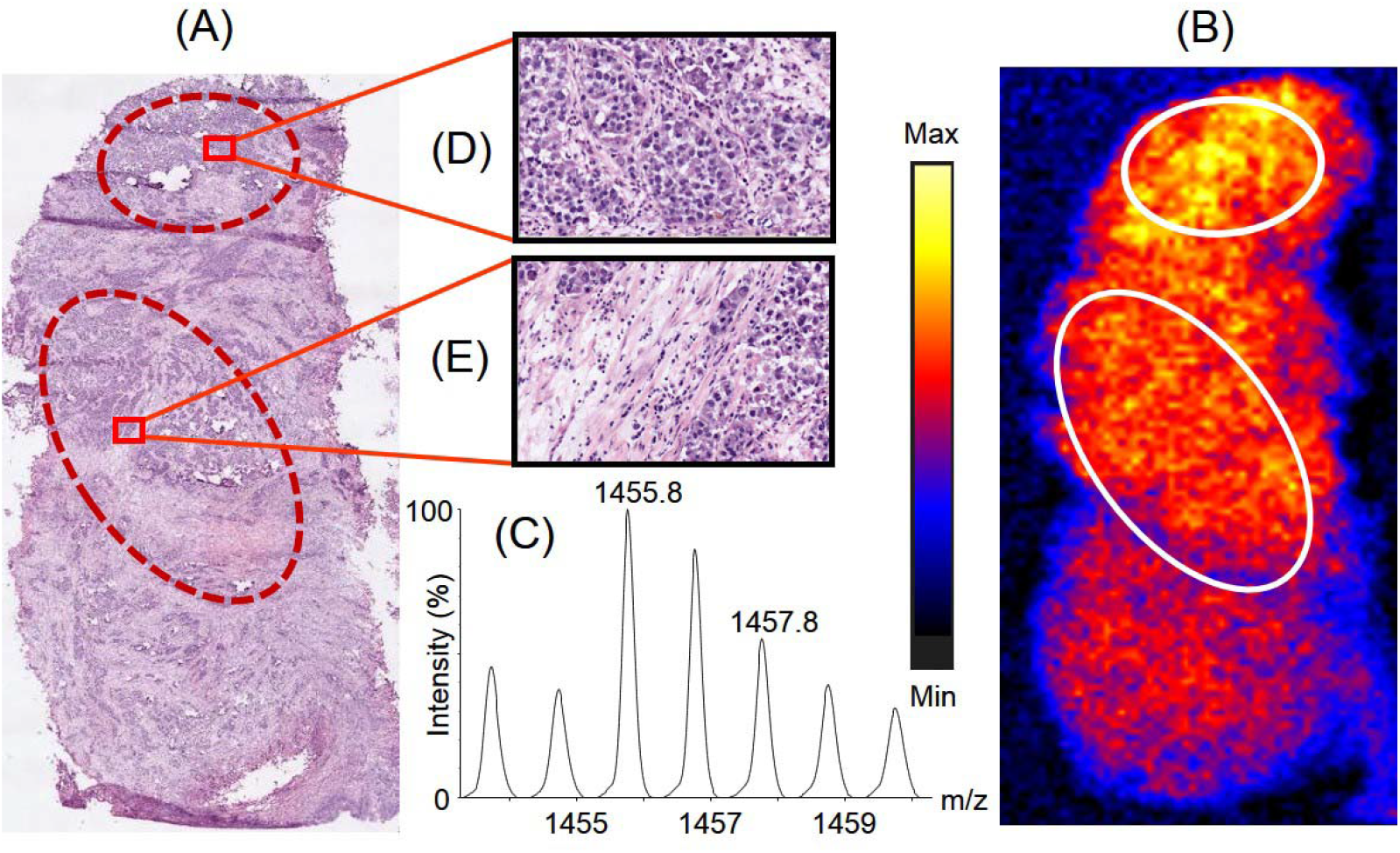
Distribution of the peptide with m/z 1455.8 across the prostate cancer tissue section and its correlation with the histopathological information. Panel A represents the H & E stained serial section of the tissue section which was used for histopathological examination. Panel B represents the distribution of the peptide across the tissue section, observed upon imaging the tissue section using MALDI mass spectrometry based imaging platform. The encircled parts represent different regions on the tissue section showing similar signal intensities of the peptide 1455.8. Panel C shows the isotopic distribution of the tryptic peptide in the mass spectra. Panels D and E show a representative 20X views consisting of different cellular population of a small part in the selected regions of the H & E stained tissue section corresponding to the regions marked in the MALDI tissue image

Figure 4 represents the distribution of the tryptic peptide with m/z 1175.6 across the prostate cancer tissue section. Using tissue proteomics, we observed the tryptic fragment is a part of the protein vinculin, consisting of residues ^237^MSAEINEIIR^246^. Figure 4A shows the corresponding region in the H & E stained tissue section (encircled) where vinculin was overexpressed in the MALDI image of a serial section of the same tissue, depicted in Figure 4B. The isotopic distribution of the peptide with m/z 1175.6 in the mass spectra is shown in Figure 4C. Figure 4D shows the 20X expanded view of a small representative part in the encircled region of the H & E stained tissue section, that shows vinculin being overexpressed in a region of the tissue section which was densely populated by cancerous epithelial cells that might have undergone clonal expansion (a process of acquiring clones with variable types of cancer-associated genetic mutations in addition to the initial genetic mutation which developed the cancer) and would potentially be presented with increased aggressiveness of cancer. Vinculin is coded by the VCL gene [18]. The Cancer Genome Atlas (TCGA) report showed that the VCL gene was altered and the corresponding mRNA was overexpressed in 4% of the patients screened with prostate cancer [19]. Vinculin is a cytoskeletal protein localized in the cytoplasmic face of the junctions between cells and extracellular matrix [20]. It binds to another cytoskeletal protein, talin, and potentially links the integrin proteins of the cell membranes to extracellular matrix [21]. It has been reported that the presence of vinculin in focal adhesions (protein assemblies forming the sites of contacts of cells to the extracellular matrix) is critical for integrin-mediated cell migration [21]. Previously, immunohistochemical studies on tissues with amplified VCL gene expression from patients with castration-resistant prostate cancer revealed overexpression of vinculin and it was reported that vinculin is a major driving gene of the 10q22 amplification in prostate cancer. Thus, overexpression of vinculin might be associated with enhanced tumour cell proliferation in prostate cancer pathogenesis [22]. Our data showing the overexpression of vinculin by cancerous epithelial cells of the prostate tissue is in agreement with established role for this protein in prostate cancer.

**Figure 4:**
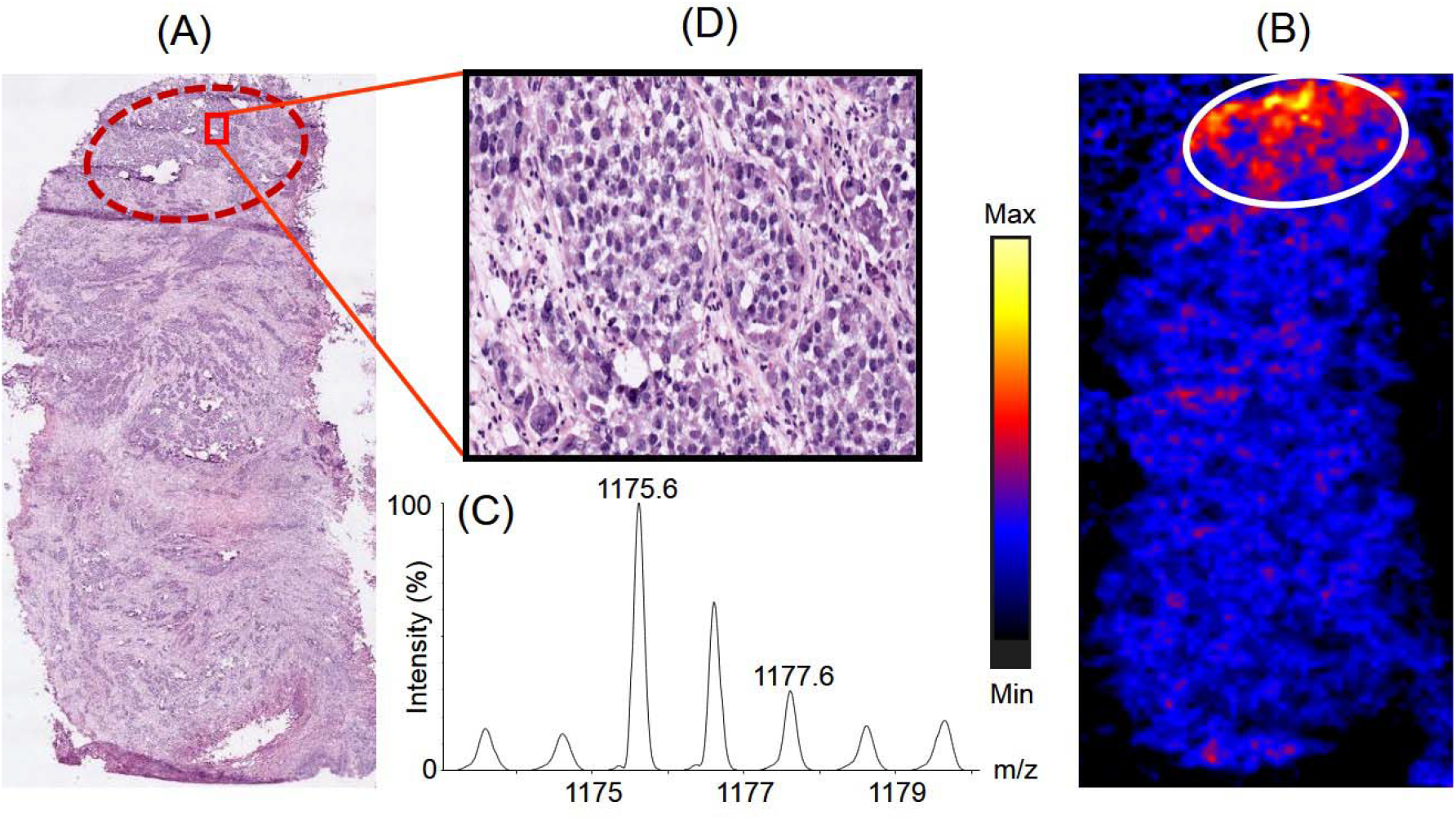
Distribution of the peptide with m/z 1175.6 across the prostate cancer tissue section and its correlation with histopathology. Panel A represents the H & E stained section of the tissue section which was used for histopathological examination. Panel B represents the distribution of the peptide across the tissue section, observed upon imaging the tissues using MALDI mass spectrometry based imaging platform. The encircled part represents the region where the intensity of the peptide was observed to be higher compared to the rest part of the tissue. Panel C shows the isotopic distribution of the tryptic peptide in the mass spectra. Panel D shows a representative 20X view the cellular population in a small part of a region on the H & E stained tissue section corresponding to the region marked in the MALDI image of the tissue having differential spatial expression of the peptide

Figure 5 represents the distribution of a tryptic peptide with m/z 1143.6 on the prostate cancer tissue section. Proteomics data revealed the peptide is a fragment of the protein, ribonuclease T2 comprising of residues ^144^ELDLNSVLLK^153^. Figure 5A shows the corresponding region in the H & E stained tissue section (encircled) where ribonuclease T2 was overexpressed in the MALDI image of a serial section of the same tissue, depicted in Figure 5B. The isotopic distribution of the peptide with m/z 1143.6 in the mass spectra is shown in Figure 5C. Figure 5D shows the 20X expanded view of a small representative part in the encircled region of the H & E stained tissue section illustrating the cellular population in this region. Upon correlation with the histopathological image, the MALDI image of the distribution of ribonuclease T2 across the tissue section showed that it was overexpressed in a region infiltrated with immune cells such as the macrophages and neutrophils. Ribonuclease T2, coded by the human gene RNASET2, is secreted by epithelial cells into the extracellular milieu [23]. TCGA database reported an alteration of RNASET2 gene and a subsequent mRNA overexpression in 4% of patients with prostate cancer [19]. The gene was classified as belonging to the category of tumor antagonizing genes/malignancy suppressing genes (TAG/MSG) [24]. TAG/MSG class of genes are known to code for proteins which are required to mediate cellular responses in the tumor microenvironment to prevent the processes that lead to malignancy, such as angiogenesis. It has been reported that overexpression of RNASET2 gene results in a triggered response from immune cells such as the monocytes and macrophages, indicating a biological mechanism based on microenvironmental control to prevent growth of tumors [24]. In addition, the activity of the protein, antigen target A kinase anchor protein (AKAP4), involved in several malignant properties of tumor, such as arresting cell cycle and preventing apoptosis in several cancers like prostate cancer, esophageal cancer, multiple myeloma, etc [25–28], is actively suppressed by ribonuclease T2 [29]. Amphiregulin, another protein involved in regulation of cancer cell invasiveness, is reported to be negatively regulated by ribonuclease T2 [30]. Therefore, in the present study, our observations on the overexpressed of Ribonuclease T2 in a region infiltrated with macrophages and neutrophils corroborated well with the literature information available for this tumor suppressor protein.

**Figure 5:**
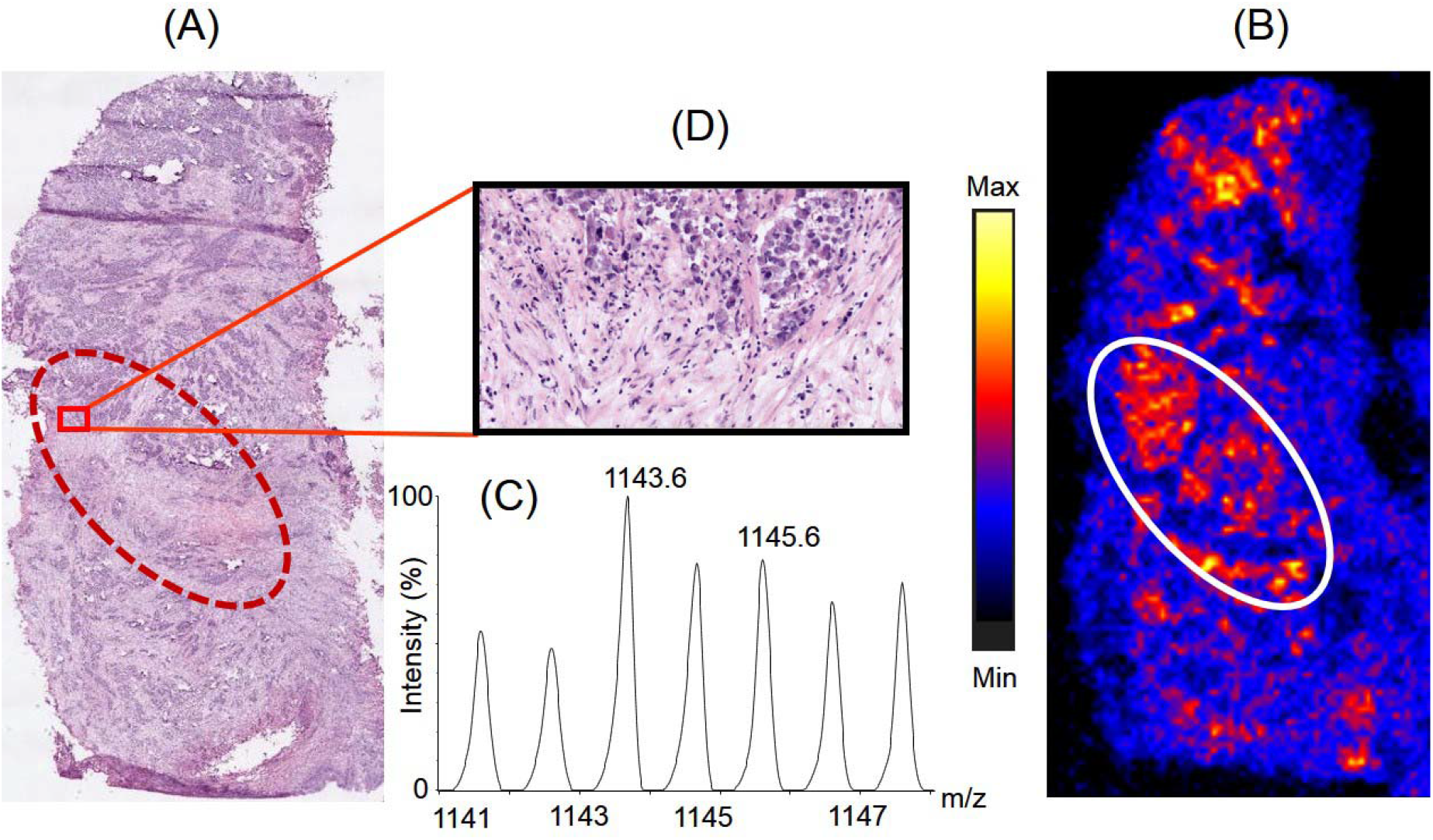
Distribution of the peptide with m/z 1143.6 across the prostate cancer tissue section and its correlation with histopathology. Panel A represents the H & E stained section of the tissue section which was used for histopathological examination. Panel B represents the distribution of the peptide across the tissue section, observed upon imaging the tissues using MALDI mass spectrometry based imaging platform. The encircled part represents the region where the intensity of the peptide was observed to be higher compared to the rest part of the tissue. Panel C shows the isotopic distribution of the tryptic peptide in the mass spectra. Panel D shows a representative 20X view the cellular population in a small part of the region on the H & E stained tissue section corresponding to the region on the tissue image having differential spatial expression the peptide

Figure 6 illustrates the distribution of a tryptic peptide with m/z 2295.2 on the prostate cancer tissue section. Proteomics data revealed the peptide is a fragment of 60 kDa heat shock protein comprising of residues ^371^IQEIIEQLDVTTSEYEKEK^389^. Figure 6A shows the corresponding region in the H & E stained tissue section (encircled) where 60 kDa heat shock protein was overexpressed in the MALDI image of a serial section of the same tissue, depicted in Figure 6B. The isotopic distribution of the peptide with m/z 2295.2 in the mass spectra is shown in Figure 6C. Figure 6D shows the 20X expanded view of a small representative part in the encircled region of the H & E stained tissue section providing insights into the cellular population in this region. In our study, we observed that 60 kDa heat shock protein was overexpressed in regions of the tissue having cancerous epithelial cells surrounded by variable amounts of stroma containing immune cells. 60 kDa heat shock protein is coded by the gene HSP60 and belongs to the family of chaperonins. It is present in the cytosol, cell surface, and extracellular space and in peripheral blood [31]. However, it is most abundantly located in the eukaryotic cell mitochondria and are induced in response to stresses, such as heat shock [32]. It has important functions in transport and folding of mitochondrial proteins. This protein is also reported to play a role in transformation, promotion of angiogenesis, and metastasis of cancer cells [33,34]. According to the TCGA report, the mRNA of the 60 kDa heat shock protein gene is overexpressed in 7% of the patients with prostate cancer [19]. An increased expression of the gene has been reported in multiple cancers such as glioblastoma, colorectal cancer, gastric cancer, breast cancer, ovarian cancer, liver cancer, bladder cancer and head and neck cancer [35]. In the present study, 60 kDa heat shock protein was found to be localized in different regions of the tissue sections having well-organized patterns of cancerous epithelial cells surrounded by variable amounts of stroma. This might indicate that 60 kDa heat shock protein was secreted in the extracellular region by the cancerous epithelial cells of the prostate gland. Therefore, our observations of the specific distribution of 60 kDa heat shock protein in prostate cancer tissues corroborated well with the reported literature information about this protein.

**Figure 6:**
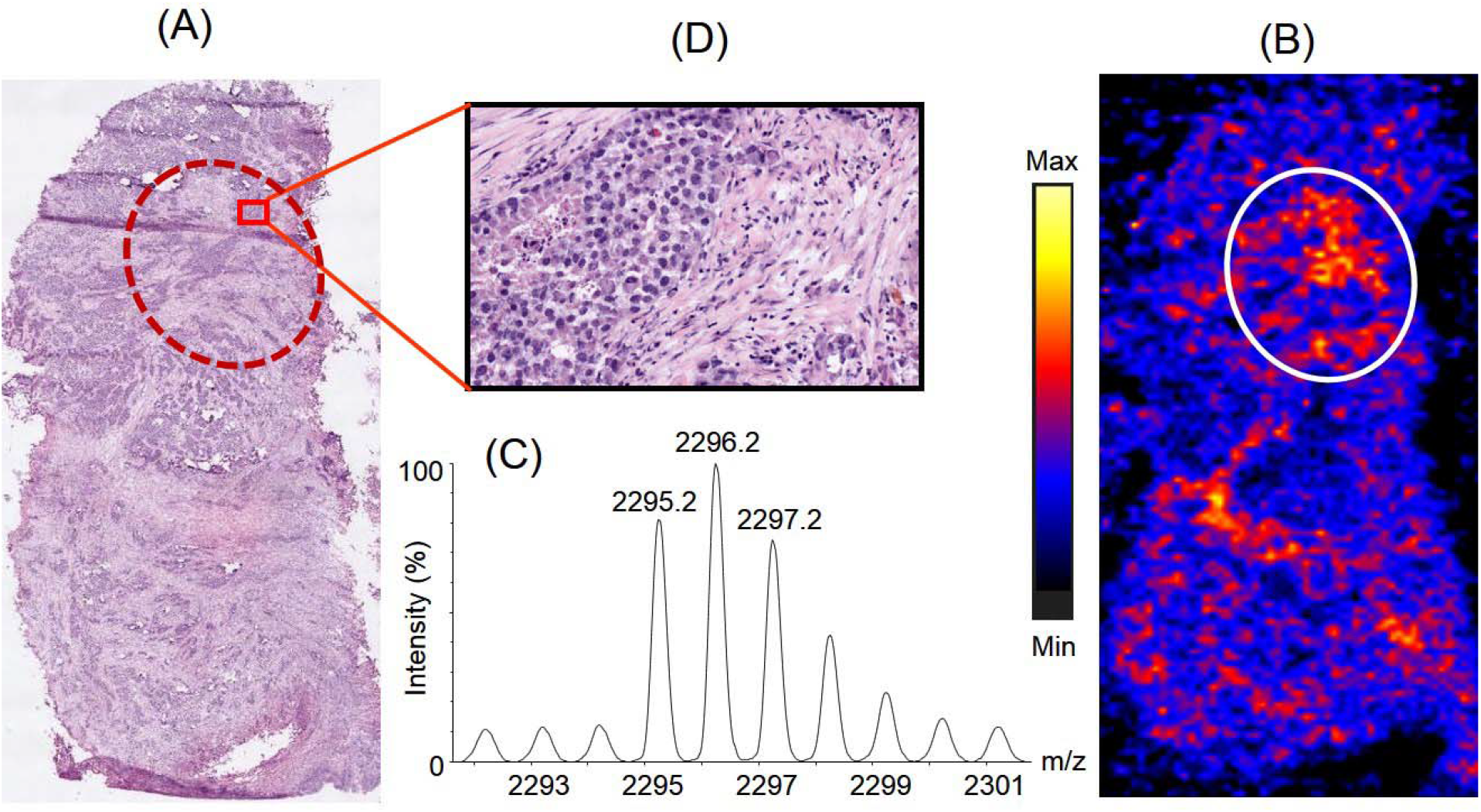
Distribution of the peptide with m/z 2295.2 across the prostate cancer tissue section and its correlation with histopathology. Panel A represents the H & E stained section of the tissue section which was used for histopathological examination. Panel B represents the distribution of the peptide across the tissue section, observed upon imaging the tissues using MALDI mass spectrometry based imaging platform. The encircled parts represent the regions where the peptide was observed to be higher expression compared to the rest part of the tissue. Panel C shows the isotopic distribution of the tryptic peptide in the mass spectra. Panel D show a representative 20X view the cellular population in a small part of the region on the H & E stained tissue section corresponding to the region on the tissue image having differential spatial expression of the peptide

We also performed MALDI IMS analysis of si needle biopsies. Four representative images of distribution of these three peptides with m/z 1175.6 of vinculin, 1143.6 of ribonuclease T2 and 2295.2 of 60 kDa heat shock protein in four representative needle biopsies collected from patients diagnosed with prostate cancer are shown in Figure S1.

In addition, we performed the quantitative proteomics analysis of the tissue samples collected from patients with prostate cancer and compared the proteome profiles with that obtained from patients with BPH, considered as control in the present study. Ribonuclease T2 and 60 kDa heat shock protein were observed to be overexpressed by 22.26- and 6 folds respectively in the prostate cancer tissue proteomes (Table 1), which corroborated well with the observations of IMS study. However, compared to BPH samples, there was no significant change in the expression of vinculin in prostate cancer tissues (0.96 folds). The proteomics data was normalized by dividing the amount of each protein by the summation of the amounts of all proteins present in the sample. The experimental setup to perform tissue proteomics includes homogenization of the tissues, which results in a loss of the heterogeneous spatial distribution of the proteins across the tissue section and disappearance of the concentration gradient of the marker protein in the pathologically important portions of the experimental tissue section with respect to its total protein content. This might be the reason that we were unable to observe the differential expression levels of vinculin, which was observed to be specifically located in a small area in the MALDI image of the tissue having dense population of cancer cells.

**Table 1:** Comparison of the relative expression of vinculin, ribonuclease T2 and 60 kDa heat shock protein between proteomes in BPH and prostate cancer tissues.

| **Accession No.** | **Protein** | **BPH**<br>**(normalized and averaged)** | **Prostate cancer**<br>**(normalized and averaged)** | **Fold change in expression**<br>**(Sample/Control)** |
| --- | --- | --- | --- | --- |
| **P18206** | Vinculin | 0.0022 | 0.00211 | 0.96 |
| **O00584** | Ribonuclease T2 | 0.00002 | 0.00041 | 22.26 |
| **P10809** | 60 kDa heat shock protein | 0.0003 | 0.0018 | 6 |

In the current scenario, histopathology is used as the gold-standard technique in the routine diagnosis of tissue sections. However, as reported, they might not be able to detect cancer in tissues at very early stages when the morphological changes in the cells are minimal [36,37]. Analyzing the changes in protein profiles in the diagnostic pathway of the disease might be additive to the currently used modalities. Cancerous tissues are marked by a change in the distribution of molecules which might arise as a result of differential regulation of the gene expression or of proteins and their post-translational modifications [38–40]. In this context, mass spectrometry based imaging approach might significantly contribute to explore the pathogenesis of the disease and also in the discovery of the disease biomarkers in future.

In the present study, we used MALDI mass spectrometry based imaging platform to explore the distribution of protein on human prostate cancer tissue sections. We observed that three proteins, vinculin, ribonuclease T2 and 60 kDa heat shock protein, previously implicated in prostate cancer pathogenesis, were differentially expressed across a prostate tissue section with cancer. To the best of our knowledge, this is the first report where MALDI imaging platform was used to visualize the differential distribution and expression profiles of these three proteins in human prostate cancer tissue sections. This lends support for the use of this approach to be applied systematically for the proteomic analysis of cancer tissues. Such studies may help identify hitherto unrecognized proteins involved in the pathogenesis of specific cancers. The advantage of this technique is its label free approach that does not depend on specific reagents, and the retention of the two-dimensional spatial and cellular heterogeneous distribution of specific markers on tissues as *in vivo*.

## Conclusions

In the present study, MALDI mass spectrometry based tissue imaging platform was used to image the distribution of multiple proteins on human prostate tissues which were diagnosed with cancer. We observed the differential expression of three proteins, vinculin, ribonuclease T2 and 60 kDa heat shock protein across the tissue sections in the regions that were found to be pathologically significant in the diagnosis of prostate cancer. We propose that upon validation of the observed overexpression of ribonuclease T2 and 60 kDa heat shock protein in prostate cancer in a larger number of patient samples might help to develop these proteins as potential biomarkers for the diagnosis of prostate cancer in future.

## Supporting information

Supplementary doc

## Acknowledgement

We acknowledge all the volunteers for providing tissue samples for the study. We acknowledge Rajiv Gandhi University of Health Sciences, Bangalore, for funding the study and Department of Science and Technology (Nanomission), Govt. of India, for funding mass spectrometry facility at St. John’s Research Institute. We are grateful to Dr. Geetashree Mukherjee and Swarnendu Bag, Tata Medical Centre, Kolkata for microscopic scanning of the H & E stained tissue sections. We also acknowledge Prof. Debnath Pal for his valuable inputs in MALDI image analysis.

## Conflict of interests

The authors declare no conflict of interests.

## Notes

### Competing Interest Statement

The authors have declared no competing interest.

