## Supplementary doc for "Differential expression of proteins in human prostate cancer tissues probed by MALDI imaging mass spectrometry"

Amrita Mitra^1^, Surya Kant Choubey^2^, Rajdeep Das^1^, Pritilata Rout^3^, Subhamoy Chatterjee^4^, Sreekanth Reddy^4^, T. S Sridhar^5^, Deepak Mishra^6^, Dipankar Banerjee^4^, Amit Kumar Mandal^1^*

^1^Clinical Proteomics Unit, Division of Molecular Medicine, St. John's Research Institute, St. John's National Academy of Health Sciences, 100ft Road, Koramangala, Bangalore – 560034

^2^Department of Urology and Renal Transplantation, St. John's Medical College & Hospital, St. John's National Academy of Health Sciences, 100ft Road, Koramangala, Bangalore – 560034

^3^Department of Pathology, St. John's Medical College, St. John's National Academy of Health Sciences, 100ft Road, Koramangala, Bangalore – 560034

^4^Indian Institute of Astrophysics, Sarjapur Main Road, 2nd Block, Koramangala, Bangalore - 560034

^5^Breast Cancer Unit, Division of Molecular Medicine, St. John’s Research Institute, 100ft Road, Koramangala, Bangalore - 560034

^6^Department of Pathology, Tata Medical Center, New Town, Rajarhat, Kolkata - 700156

*Corresponding Author

Dr. Amit Kumar Mandal

Clinical Proteomics Unit

Division of Molecular Medicine

St. John's Research Institute,

St. John's National Academy of Health Sciences

100ft Road, Koramangala,

Bangalore – 560034, India

Present address:

Dr. Amit Kumar Mandal

Department of Biological Sciences,

Indian Institute of Science Education and Research Kolkata

Mohanpur, Nadia, West Bengal

India

Pin-741246

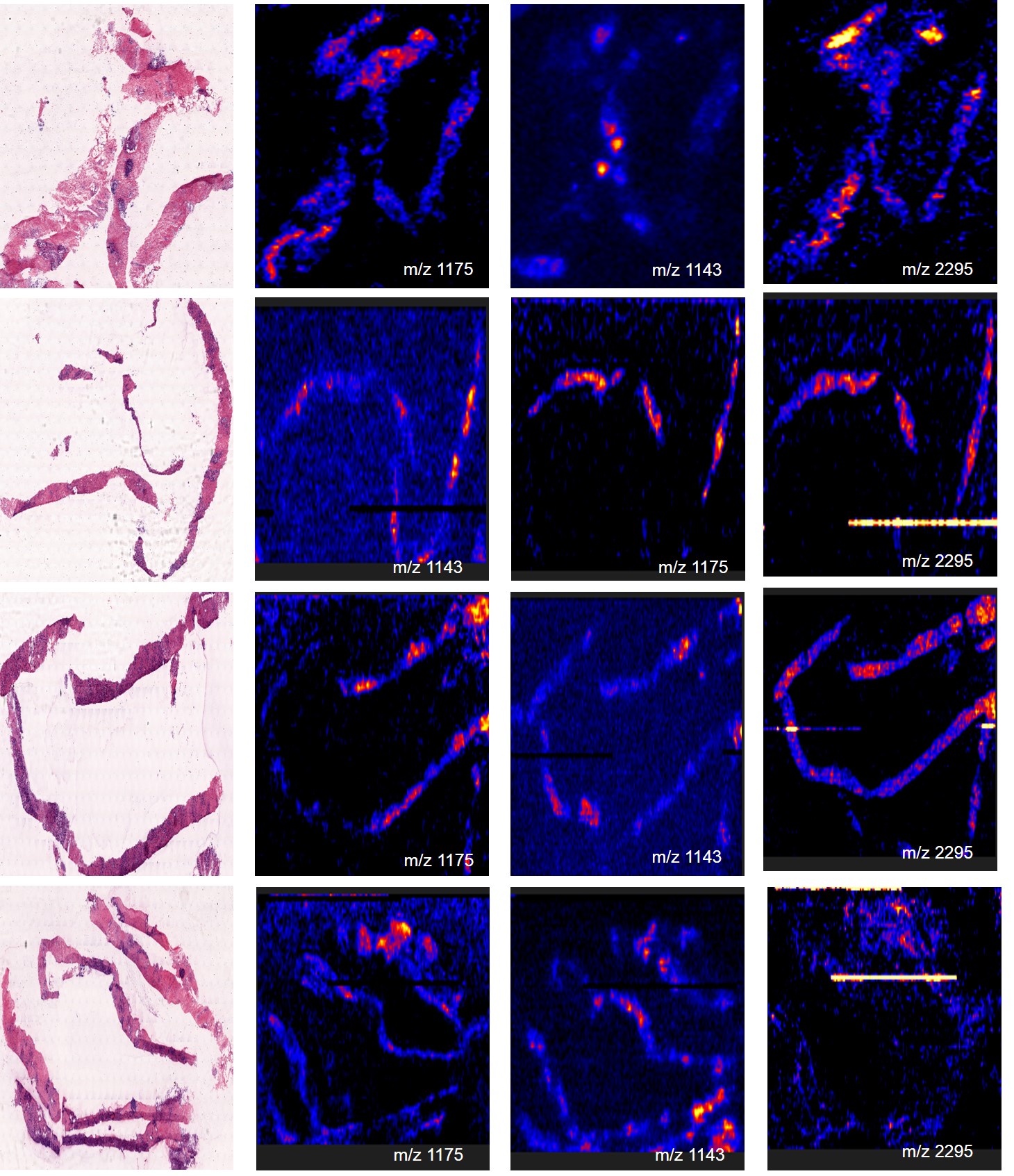

**Figure S1:** Distribution of the tryptic peptides with m/z 1175.6 of vinculin, 1143.6 of ribonuclease T2 and 2295.2 of 60 kDa heat shock protein across prostate needle core biopsy tissue sections obtained from four patients diagnosed with prostate cancer
